# decoupleR: Ensemble of computational methods to infer biological activities from omics data

**DOI:** 10.1101/2021.11.04.467271

**Authors:** Pau Badia-i-Mompel, Jesús Vélez, Jana Braunger, Celina Geiss, Daniel Dimitrov, Sophia Müller-Dott, Petr Taus, Aurelien Dugourd, Christian H. Holland, Ricardo O. Ramirez Flores, Julio Saez-Rodriguez

## Abstract

**Summary:** Many methods allow us to extract biological activities from omics data using information from prior knowledge resources, reducing the dimensionality for increased statistical power and better interpretability. Here, we present decoupleR, a Bioconductor package containing computational methods to extract these activities within a unified framework. decoupleR allows us to flexibly run any method with a given resource, including methods that leverage mode of regulation and weights of interactions. Using decoupleR, we evaluated the performance of methods on transcriptomic and phospho-proteomic perturbation experiments. Our findings suggest that simple linear models and the consensus score across methods perform better than other methods at predicting perturbed regulators.

**Availability and Implementation:** decoupleR is open source available in Bioconductor (https://www.bioconductor.org/packages/release/bioc/html/decoupleR.html). The code to reproduce the results is in Github (https://github.com/saezlab/decoupleR_manuscript) and the data in Zenodo (https://zenodo.org/record/5645208).

**Contact:** Julio Saez-Rodriguez at.

## 1. Introduction

Omics datasets, such as transcriptomics or phospho-proteomics, provide unbiased high-dimensional molecular profiles. However, their big dimensionality, combined with the highly connected nature of the molecules that are measured, make it difficult to interpret them in a mechanistically relevant manner. Leveraging prior knowledge we can use computational methods to infer which biological activities are relevant. For example, the activity of transcription factors and kinases can be inferred robustly from downstream transcripts and phosphosite targets, respectively (Dugourd and Saez-Rodriguez, 2019). Over the past decade a plethora of methods that infer biological activity have emerged, each with its own assumptions and biases.

Although comparisons and collections of these methods exist (Maciejewski, 2014; Mathur *et al*., 2018; Väremo *et al*., 2013; Alhamdoosh *et al*., 2017; Geistlinger *et al*., 2020; Yilmaz *et al*., 2021), they do not incorporate recent methodological developments, such as modeling activities based on weighted mode of regulation. Here we present decoupleR, an R package containing a collection of methods adapted for biological activity estimation in bulk, single cell and spatial omics data.

## 2. Implementation

Currently decoupleR contains 11 different methods (Fig. 1), these include popular methods such as AUCell (Aibar *et al*., 2017), fast GSEA (Sergushichev, 2016), GSVA (Hänzelmann *et al*., 2013), over representation analysis (ORA) (Fisher, 1922), univariate linear model (ULM) (Teschendorff and Wang, 2020), VIPER (Alvarez *et al*., 2016) and others (Supplementary Table 1). The inputs of decoupleR are: (1) a matrix containing molecular feature values like gene expression counts per sample and (2) a prior knowledge resource such as a collection of gene sets. The user can then choose any method alone or many simultaneously. Decoupler also provides a consensus score obtained by aggregating the different ranked scores (Kolde *et al*., 2012).

**Figure 1.**
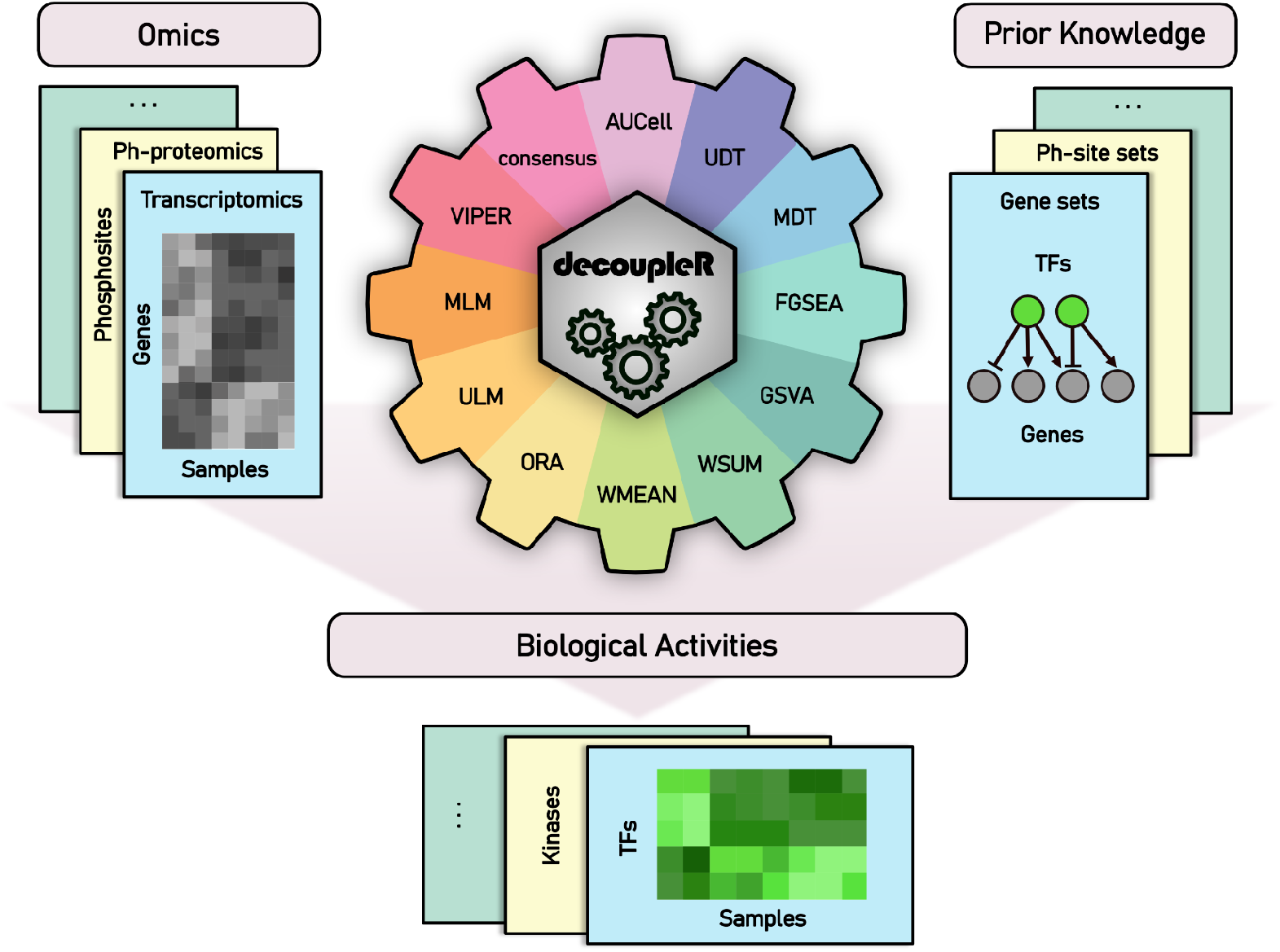
decoupleR’s workflow. decoupleR contains a collection of computational methods that coupled with prior knowledge resources estimate biological activities from omics data.

## 3. Benchmark design

We used decoupleR to evaluate the performance of individual methods by recovering perturbed regulators - transcription factors (TFs) and kinases - from two independent collections of transcriptomics (Holland *et al*., 2020) and phospho-proteomics (Hernandez-Armenta *et al*., 2017) datasets (Supplementary Note), respectively, upon single-gene perturbation experiments. As resources we used the gene regulatory network DoRothEA (Garcia-Alonso *et al*., 2019), and a kinase substrate network (Hernandez-Armenta *et al*., 2017), respectively.

We built a benchmarking pipeline with decoupleR (Supplementary Note), which evaluates the performance of regulator activity scores from different methods, mainly focused on the sensitivity of methods. Furthermore, to evaluate the robustness of the methods to noise, we added or deleted a percentage of edges from the prior knowledge resources.

## 4. Results

Methods return different distributions of activities (Supplementary Fig. S1) but display general similarities (Supplementary Fig. S2), with a median Spearman correlation of activities between methods of 0.52, and 0.66 for transcriptomics and phospho-proteomics, respectively. There was also a moderate agreement between methods in the top 5% ranked regulators (median Jaccard indexes of 0.23 and 0.24, respectively; Supplementary Fig. S2).

Despite these similarities, methods showed different performances at predicting perturbed regulators (Supplementary Fig. S3). Some of them performed consistently better than the others (Supplementary Tab. S2), the top three being: consensus, multivariate linear model (MLM) and ULM. Moreover, methods that leverage weights perform better when those are taken into account (p-value<2.2e-16; W=2.32e+10; N=2.20e+05; one-sided Wilcoxon signed-rank test) (Supplementary Fig. S4).

Deleting edges in the resource had a greater effect than adding them across methods (Supplementary Fig. S5); with a median Spearman correlation of activities to the original ones of 0.84 and 0.77 for the addition and deletion, respectively (p-value<2.2e-16; W=1.11e+05; N=5.10e+02; one-sided Wilcoxon signed-rank test). Additionally, adding or deleting edges decreased predictability, and deleting edges had a worse effect than adding (adjusted p-values<2.2e-16 for normal-addition, <2.2e-16 for normal-deletion and 7.1e-06 for deletion-addition; F=130.68; Tukey’s HSD post-hoc test) (Supplementary Fig. S6).

Finally, we evaluated decoupleR’s speed and found that top performer methods run relatively fast (9.95e-04 seconds per sample and regulator with an Intel(R) Core(TM) i7-8550U CPU @ 1.80GHz) (Supplementary Fig. S7), enabling their use with larger datasets such as single-cell or spatial omics.

## 5. Conclusion

In summary, decoupleR combines a variety of methods to infer biological activities into one efficient, robust, and user-friendly tool. With a common syntax for different methods, types of omics datasets, and knowledge sources, it facilitates the exploration of different approaches and can be integrated in many workflows.

We observed that the majority of methods return adequate estimates of regulator activities, but that their aggregation into a consensus score and linear models perform better than other methods. We welcome the addition of further methods by the community.

## 8. Acknowledgments

We thank Celia Lerma-Martin for the design of the main figure and Attila Gabor for the technical support.

## Supplementary Information

### Methods overview

#### AUCell

AUCell (Aibar *et al*., 2017) uses the Area Under the Curve (AUC) to calculate whether a set of targets is enriched within the molecular readouts of each sample. To do so, AUCell first ranks the molecular features of each sample from highest to lowest value, resolving ties randomly. Then, an AUC can be calculated using by default the top 5% molecular features in the ranking. Therefore, this metric represents the proportion of abundant molecular features in the target set, and their relative abundance value compared to the other features within the sample.

#### Univariate Decision Tree

Univariate Decision Tree (UDT) (Therneau and Atkinson, 2019) fits a single decision tree for each regulator and sample. As a unique covariable, UDT uses the associated weights of a given regulator to estimate the molecular readouts of all molecular features in a sample. Target features with no associated weight are set to zero. The obtained feature importance from the fitted model is the activity of the regulator.

#### Multivariate Decision Trees

Multivariate Decision Trees (MDT) (Wright and Ziegler, 2017) fits an ensemble of decision trees, known as random forest, to infer regulator activities. MDT, contrary to UDT, uses all regulators of a given network to estimate the molecular readouts of all molecular features in a sample. Same as UDT, target features with no associated weight are set to zero. The feature importances extracted from the fitted model are the regulator activities.

#### Fast Gene Set Enrichment Analysis

Fast Gene Set Enrichment Analysis (FGSEA) (Sergushichev, 2016) estimates regulator activities using a GSEA implementation based on an adaptive multi-level split Monte Carlo scheme. In GSEA, molecular features are first ranked per sample. Then, an enrichment score (ES) is calculated by walking down the list of features, increasing a running-sum statistic when a feature in the target feature set is encountered and decreasing it when it is not. The magnitude of the increment depends on the correlation of the molecular feature with the regulator being evaluated. The final ES is the maximum deviation from zero encountered in the random walk. Finally, a normalized ES (NES), called *norm_fgsea* in decoupleR, can be calculated using permutations.

#### Gene Set Variation Analysis

Gene Set Variation Analysis (GSVA) (Hänzelmann *et al*., 2013) starts by transforming the input molecular readouts matrix to a readout-level statistic using Gaussian kernel estimation of the cumulative density function. Then, readout-level statistics are ranked per sample and normalized to up-weight the two tails of the rank distribution. Afterwards, an enrichment score (ES) is calculated as in GSEA, using the running sum statistic. Finally, the ES can be normalized by subtracting the largest negative ES from the largest positive ES.

#### Weighted Sum

Weighted Sum (WSUM) infers regulator activities by first multiplying each target feature by its associated weight which then are summed to a final enrichment score (ES). It can be defined as:

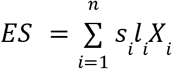

Where *n* is the number of targets for a given regulator, *s*_*i*_ is the associated mode of regulation (either positive of negative), *l*_*i*_ is the likelihood of that event happening and *X*_*i*_ is a molecular feature statistics like gene expression. In case *s*_*i*_ or *l*_*i*_ are not present, these are set to one.

Furthermore, permutations of random target features can be performed to obtain a normalized score (NES), called *norm_wsum* in decoupleR, with *R* being the obtained random null distribution:

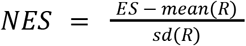

A corrected enrichment score (CES), called *corr_wsum*, is also obtained:

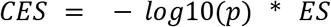

Where *p* is the empirical p-value defined as:

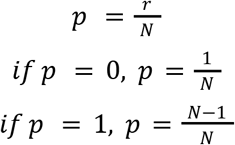

Here, *r* is the number of times *R* was bigger than the absolute value of ES and *N* is the number of random permutations. NES and CES are alike, but CES can handle better zero inflated distributions since NES requires a high *N* value to avoid having a *sd*(*R*) equal to zero.

#### Weighted Mean

Weighted Mean (WMEAN) is similar to WSUM but it divides the obtained ES by the sum of the absolute value of weights. It can be defined as:

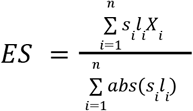

Like in WSUM, a NES (*norm_wmean*) and a CES (*corr_mean*) can be calculated if random permutations of target features are performed. It is worth mentioning that *norm_wmean* and *norm_wsum* converge into the same scores since their null distributions are the same.

#### Over Representation Analysis

Over Representation Analysis (ORA) (Fisher, 1922) measures the overlap between the target feature set and a list of most altered molecular features in the input matrix. The most altered molecular features can be selected from the top and/or bottom of the molecular readout distribution. ORA first builds a contingency table and then runs a one-tailed Fisher’s exact test to determine if a regulator’s set of features are enriched in the selected features from the data. The resulting score is:

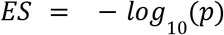

Where *p* is the obtained p-value from the test.

#### Univariate Linear Model

Univariate Linear Model (ULM) (Teschendorff and Wang, 2020), like UDT, uses as a unique covariable the weighted mode of regulation of a single regulator to estimate the molecular readouts of all molecular features in a sample. Target features with no associated weight are set to zero. The obtained t-value from the fitted model is the activity of the regulator.

#### Multivariate Linear Model

Multivariate Linear Model (MLM), contrary to ULM and similar to MDT, uses all regulators of a given network to estimate the molecular readouts of all molecular features in a sample. Same as ULM, target features with no associated weight are set to zero and the obtained t-values from the fitted model are the activities of the regulators.

#### VIPER

Virtual Inference of Protein-activity by Enriched Regulon analysis (VIPER) (Alvarez *et al*., 2016) estimates biological activities by performing a three-tailed enrichment score calculation. First, a ranking is performed for the absolute value of the molecular statistics in the input matrix per sample. The closer value to zero in the matrix is given a ranking of one and the most extreme positive value is given a ranking of N. Then, these rankings are quantile transformed. The one-tailed enrichment score is computed as:

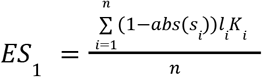

Here, *n* is the number of targets for a given regulator, *s*_*i*_ is the associated mode of regulation, *l*_*i*_ is the likelihood of that interaction and *K*_*i*_ is the quantile-transformed ranking of molecular statistics. Next, molecular targets inside each regulator are ranked again, now based on the mode of regulation, either positive or negative:

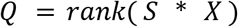

*S* is a vector indicating the mode of regulation for each target feature and *X* is a vector containing the molecular statistics from a given sample. Ranks are also quantile transformed and the two-tailed ES is calculated as:

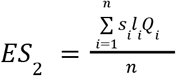

*Q*_*i*_ is the two-tailed quantile-transformed ranking of molecular statistics. Then, the three-tail score is defined as:

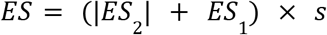

Where *s* is the sign of *ES*_2_. Finally a normalized enrichment score is estimated by:

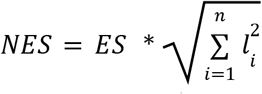

Which is an analytical approximation to random permutations.

#### Consensus

A consensus score is generated when more than one method is run with decoupleR. The score is generated using Robust Rank Aggregation (Kolde *et al*., 2012), using a probabilistic model for aggregation that is robust to noise and facilitates the calculation of significance probabilities for all the elements in the final ranking. The resulting score is:

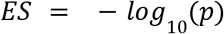

Where *p* is the obtained p-value from the model.

#### Benchmark design

We used decoupleR to evaluate the performance of individual methods by recovering perturbed transcription factors (TFs) from a curation of single-gene perturbation experiments (Holland *et al*., 2020). As a resource we used DoRothEA, a gene regulatory network linking TFs to target genes by their mode of regulation (Garcia-Alonso *et al*., 2019). Perturbation experiments where the targeted regulator was not in DoRothEA were removed. After filtering, this dataset is composed of gene expression data from 92 knockdown and overexpression experiments of 40 unique TFs in human cells. Additionally, we tested the performance of decoupleR on phospho-proteomic data. For this, we filtered in a similar fashion a curated set of knockdown and overexpression single-kinase perturbation experiments, obtaining 63 experiments including 14 unique kinases, and applied a weighted resource from the same publication that links kinases to their target phosphosites (Hernandez-Armenta et al., 2017). For the transcriptomic dataset, differential expression analysis was performed with limma (Ritchie *et al*., 2015) and the resulting t-values were used as input. For the phospho-proteomics, the quantile-normalized log2-fold changes from different studies were used to make them comparable. The unprocessed data can be accessed through Zenodo: https://zenodo.org/record/5645208.

We built a benchmarking package using decoupleR, called decoupleRBench (https://github.com/saezlab/decoupleRBench) which evaluates the performance of TF and kinase activity scores from different methods. Regulator activities were inferred from perturbation experiment data for both omics datasets using every method with default parameters. Since we only have one perturbed regulator for each experiment, we decided to concatenate all experiments into a single vector to have more than one True Positive case. Afterwards, we transformed the obtained scores to their absolute value. Since there are overexpression and knockout perturbation experiments, we assumed that perturbed regulators can have either highly positive or highly negative scores. Moreover, given that the true positive classes are limited by the TFs or kinases covered in the perturbation experiments, we added a downsampling strategy, where for each permutation an equal number of negative classes was randomly sampled. Finally, the area under the Receiver operating characteristic (AUROC) and Precision Recall curve metrics (AUPRC) were computed for each downsampling permutation. For the phospho-proteomics dataset, we ran two versions of the prior knowledge resource, one without weights and one with weights coming from kinase binding potentials, to assess whether the addition of weights gave any additional value to the prediction precision.

The obtained activities were further compared by computing the Spearman correlation between the concatenated scores of all samples from one method to another. We also checked the overlap of regulators with high absolute value score between methods by computing the Jaccard index of each pair of experiments. The final Jaccard index comes from calculating the median across experiments.

Furthermore, to evaluate the robustness of the methods to noise, we added or deleted a percentage of edges (25%, 50% and 75%) to every regulator in the prior knowledge networks. When random edges were added, their mode of regulation and weight were set to 1. For every mode (addition or deletion) and percentage, we generated five different networks, which we ran through the benchmarking pipeline of decoupleRBench. With the inferred regulator scores, for every percentage and mode we measured robustness as the correlation of scores with the normal ones and the difference of performance in AUROC and AUPRC to the normal networks.

## Supplementary tables

**Supplementary Table 1.** List of methods currently available in decoupleR. Methods are classified by whether they model the mode of regulation (Sign) or the likelihood of the source-target link (Weight), whether they are based on permutations, whether they generate a p-value associated with the inferred score and by their range of values.

| Name | Citation | Sign | Weight | Permutation | p-value | Range |
| --- | --- | --- | --- | --- | --- | --- |
| AUCell | (Aibar *et al.*, 2017) | No | No | No | No | 0,1 |
| UDT | (Therneau and Atkinson, 2019) | Yes | Yes | No | No | 0, Inf |
| MDT | (Wright and Ziegler, 2017) | Yes | Yes | Yes | No | 0, Inf |
| FGSEA | (Sergushichev, 2016) | No | No | Yes | Yes | -1, +1 |
| GSVA | (Hänzelmann *et al.*, 2013) | No | No | No | No | -1, +1 |
| WSUM | - | Yes | Yes | Yes | Yes | -Inf, +Inf |
| WMEAN | - | Yes | Yes | Yes | Yes | -Inf, +Inf |
| ORA | (Fisher, 1922) | No | No | No | Yes | 0, Inf |
| ULM | (Teschendorff and Wang, 2020) | Yes | Yes | No | Yes | -Inf, +Inf |
| MLM | - | Yes | Yes | No | Yes | -Inf, +Inf |
| VIPER | (Alvarez *et al.*, 2016) | Yes | Yes | Yes | Yes | -Inf, +Inf |
| Consensus | (Kolde *et al.*, 2012) | No | No | No | Yes | 0, Inf |

**Supplementary Table 2.** List of methods ranked by their performance in the benchmarking pipeline. Methods are ranked by the median area under the curve (AUC) of the joint distribution of all downsampling permutations in both AUROCs and AUPRCs for both datasets. Methods with significant p-values have a greater distribution of AUCs than the rest, computed using the one-sided Mann-Whitney U test (N=6.40e+05).

| Method | p-value | W | Median AUC |
| --- | --- | --- | --- |
| consensus | <2.2e-16 | 1.70e+10 | 0.68 |
| mlm | <2.2e-16 | 1.69e+10 | 0.67 |
| ulm | <2.2e-16 | 1.52e+10 | 0.66 |
| norm_wmean/norm_wsum | <2.2e-16 | 1.43e+10 | 0.65 |
| ora | <2.2e-16 | 1.42e+10 | 0.64 |
| corr_wsum | <2.2e-16 | 1.28e+10 | 0.64 |
| udt | <2.2e-16 | 1.26e+10 | 0.64 |
| mdt | 5.76e-5 | 1.21e+10 | 0.63 |
| wsum | 0.144 | 1.21e+10 | 0.64 |
| viper | 1 | 1.16e+10 | 0.62 |
| aucell | 1 | 1.13e+10 | 0.62 |
| corr_wmean | 1 | 1.09e+10 | 0.62 |
| wmean | 1 | 9.51e+09 | 0.60 |
| fgsea | 1 | 9.01e+09 | 0.59 |
| norm_fgsea | 1 | 6.86e+09 | 0.58 |
| gsva | 1 | 5.62e+09 | 0.56 |

## Supplementary figures

**Supplementary Figure 1.**
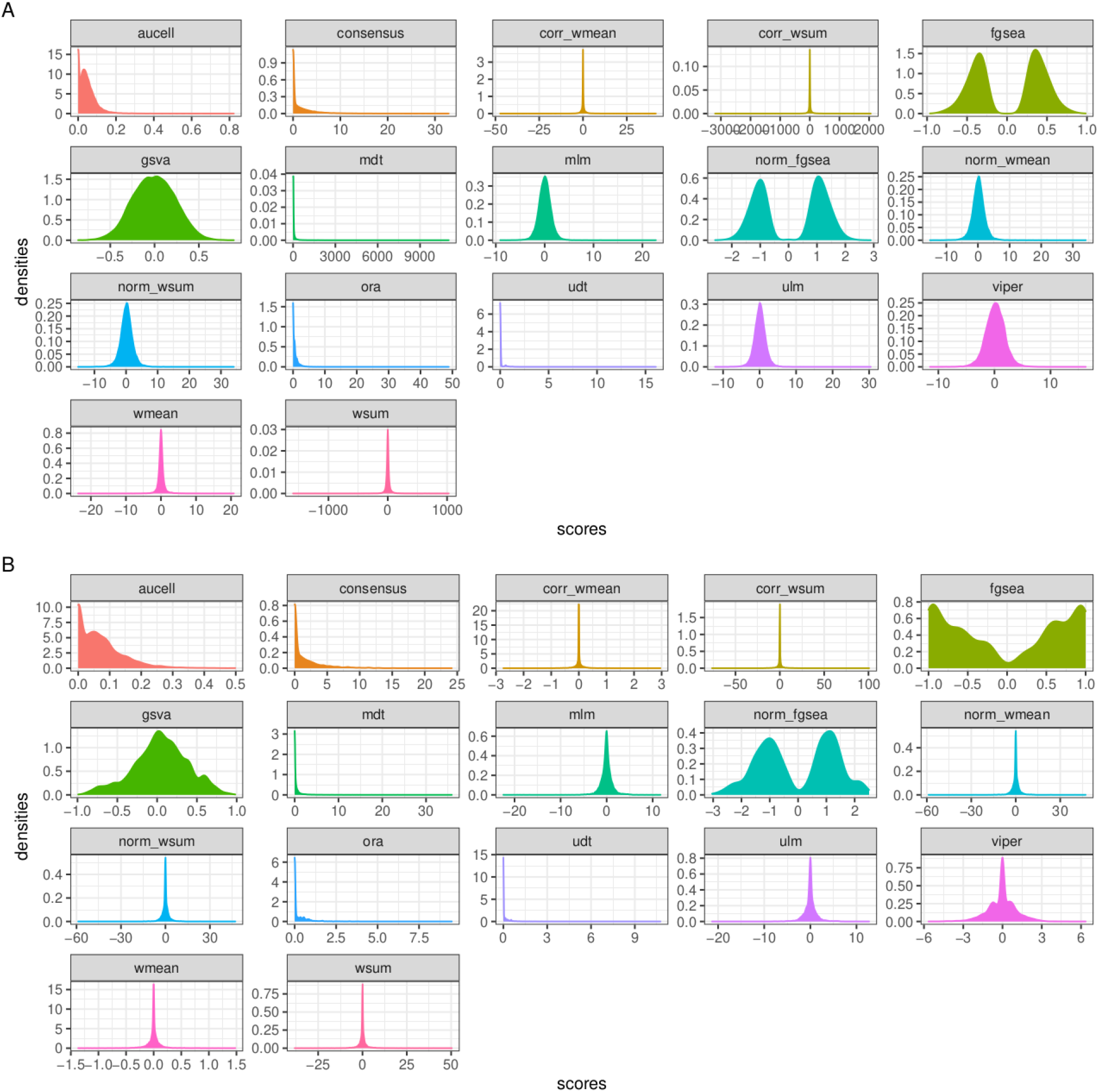
Method scores distributions for the transcriptomic dataset (A) and phospho-proteomics dataset (B).

**Supplementary Figure 2.**
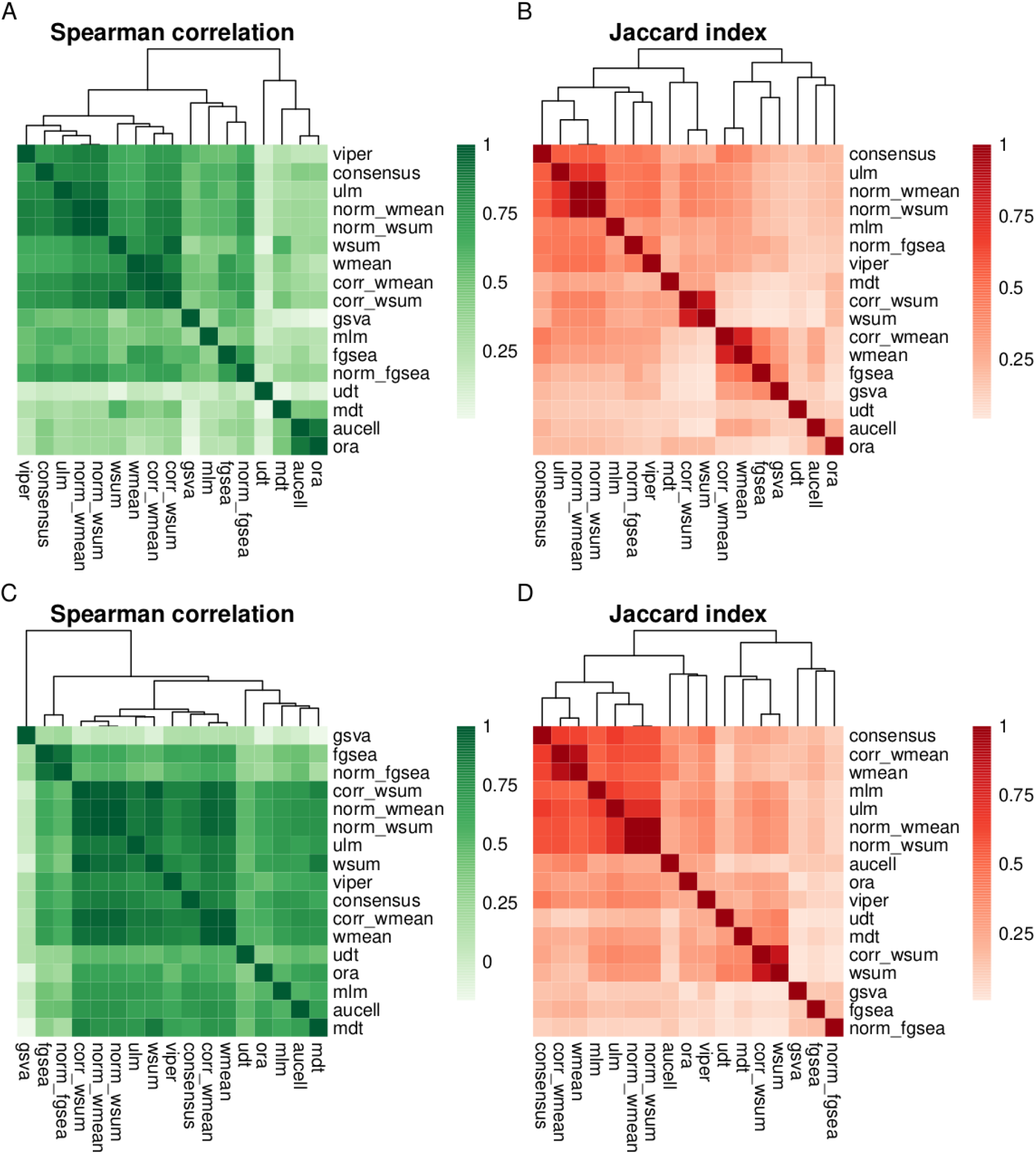
Spearman correlations between methods using the transcriptomics (A) and phospho-proteomics (C) datasets. Median Jaccard index between methods of the top 5% TFs (B) or kinases (D) ranked by the absolute value of enrichment score.

**Supplementary Figure 3.**
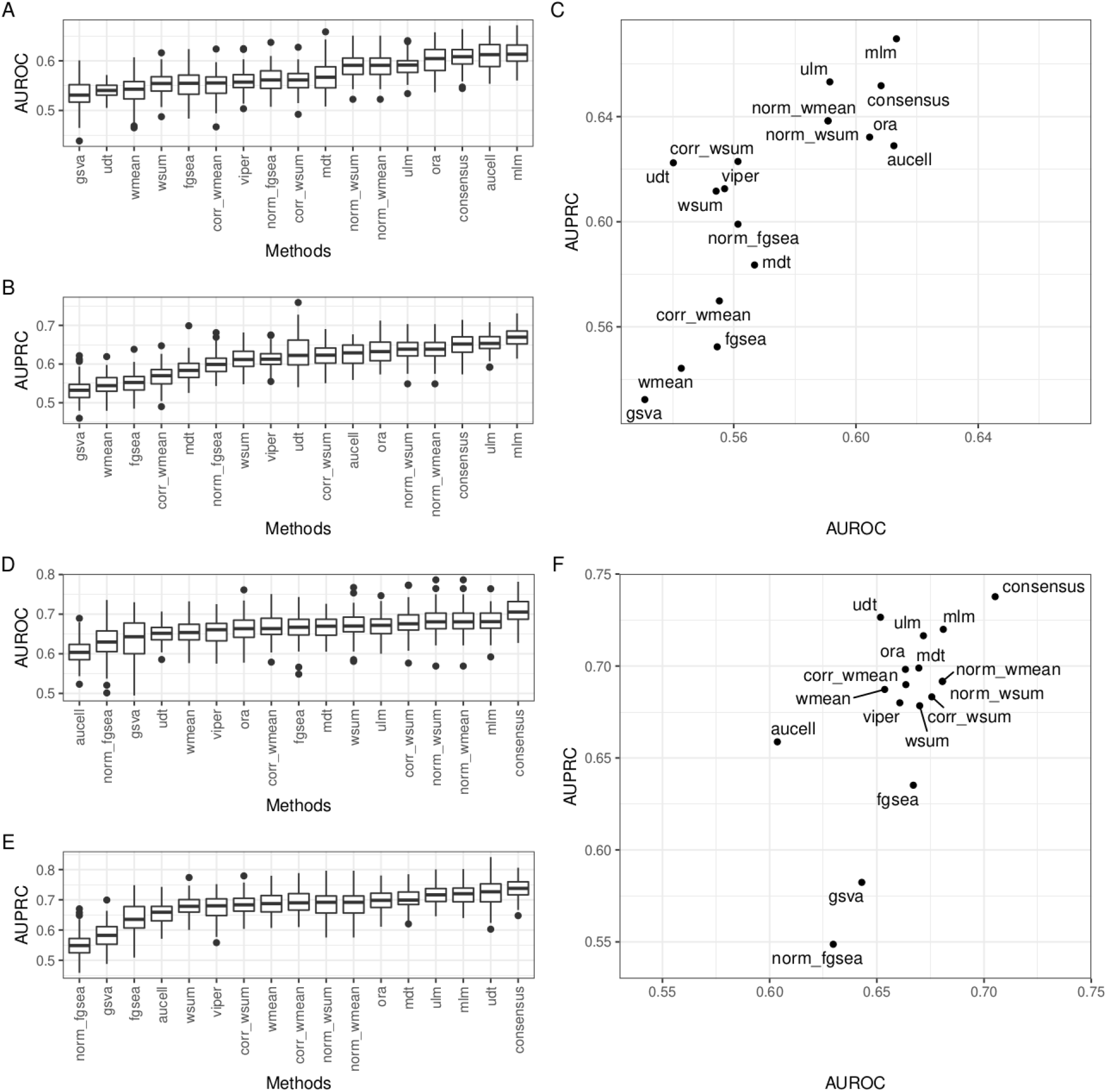
Distributions of AUROCs (A), AUPRCs (B) and the median for both (C) for each method in the transcriptomics dataset. Distributions of AUROCs (D), AUPRCs (E) and the median for both (F) for each method in the phospho-proteomics dataset.

**Supplementary Figure 4.**
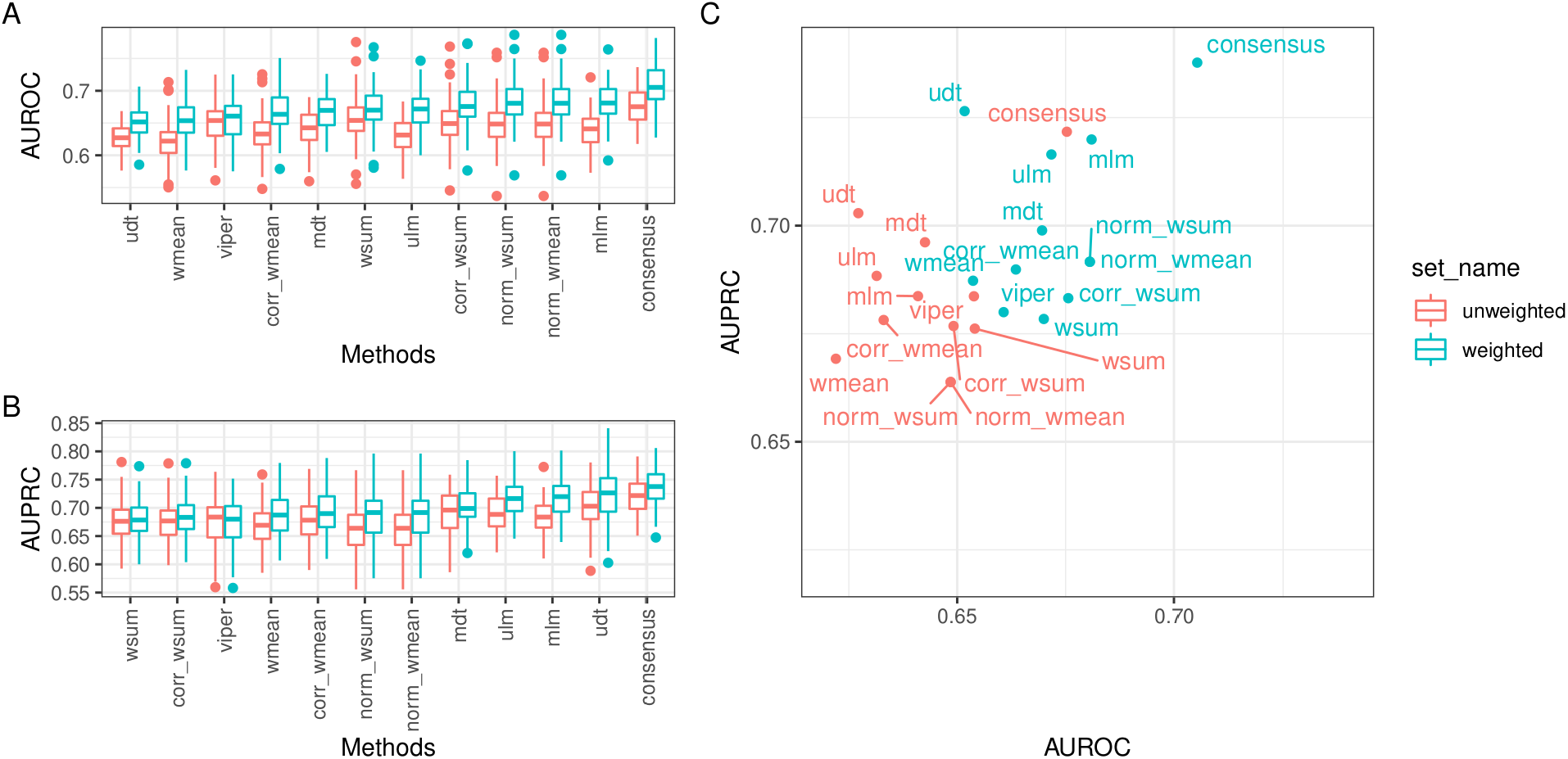
Distributions of AUROCs (A), AUPRCs (B) and the median for both (C) for each method in the phospho-proteomics dataset. Color indicates if the weights of the prior knowledge resource were used.

**Supplementary Figure 5.**
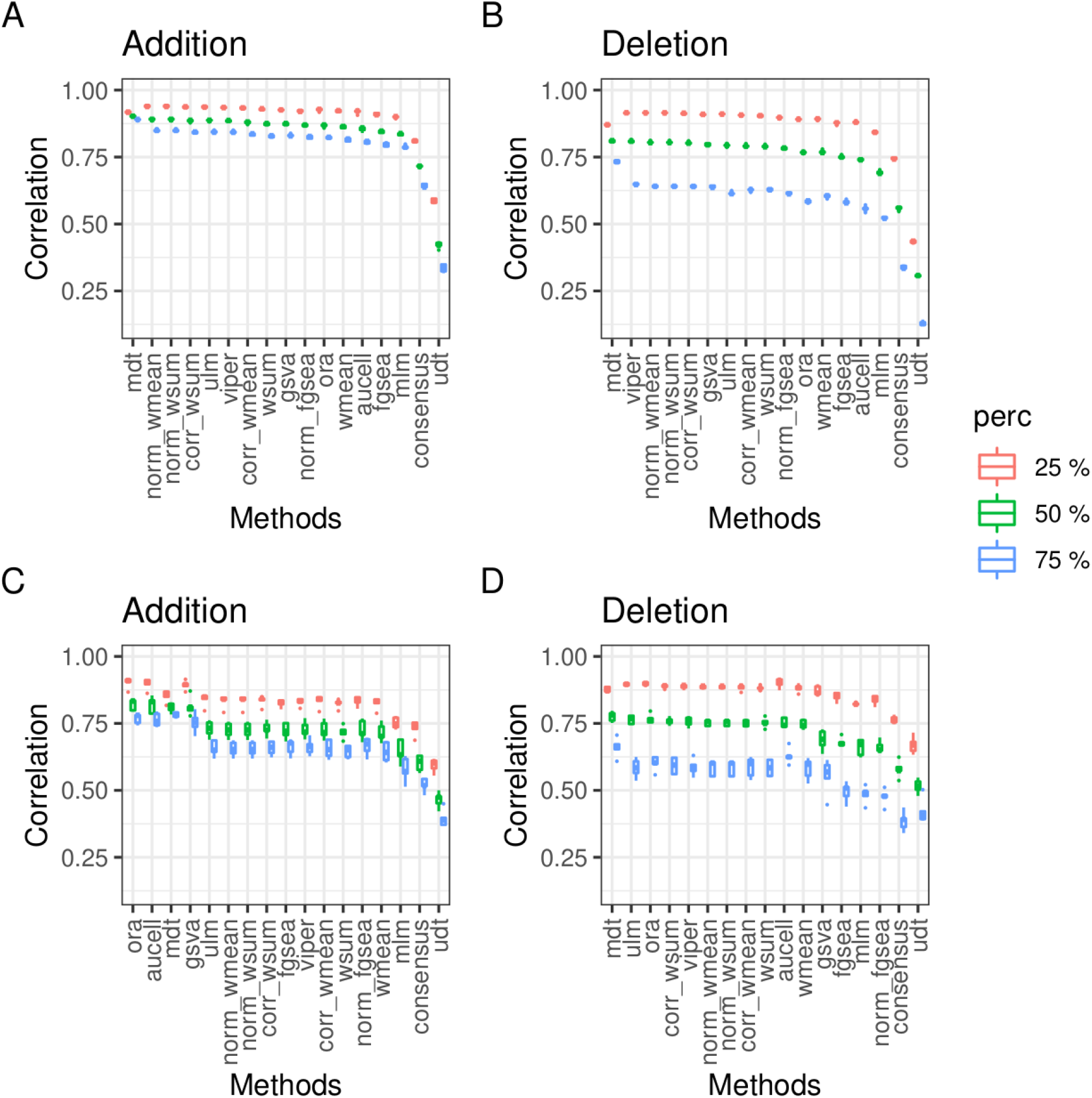
Correlations between original enrichment scores and scores obtained after adding or deleting a percentage of edges to the prior knowledge resource used for the transcriptomic (A,B) and phospho-proteomic (C,D) datasets.

**Supplementary Figure 6.**
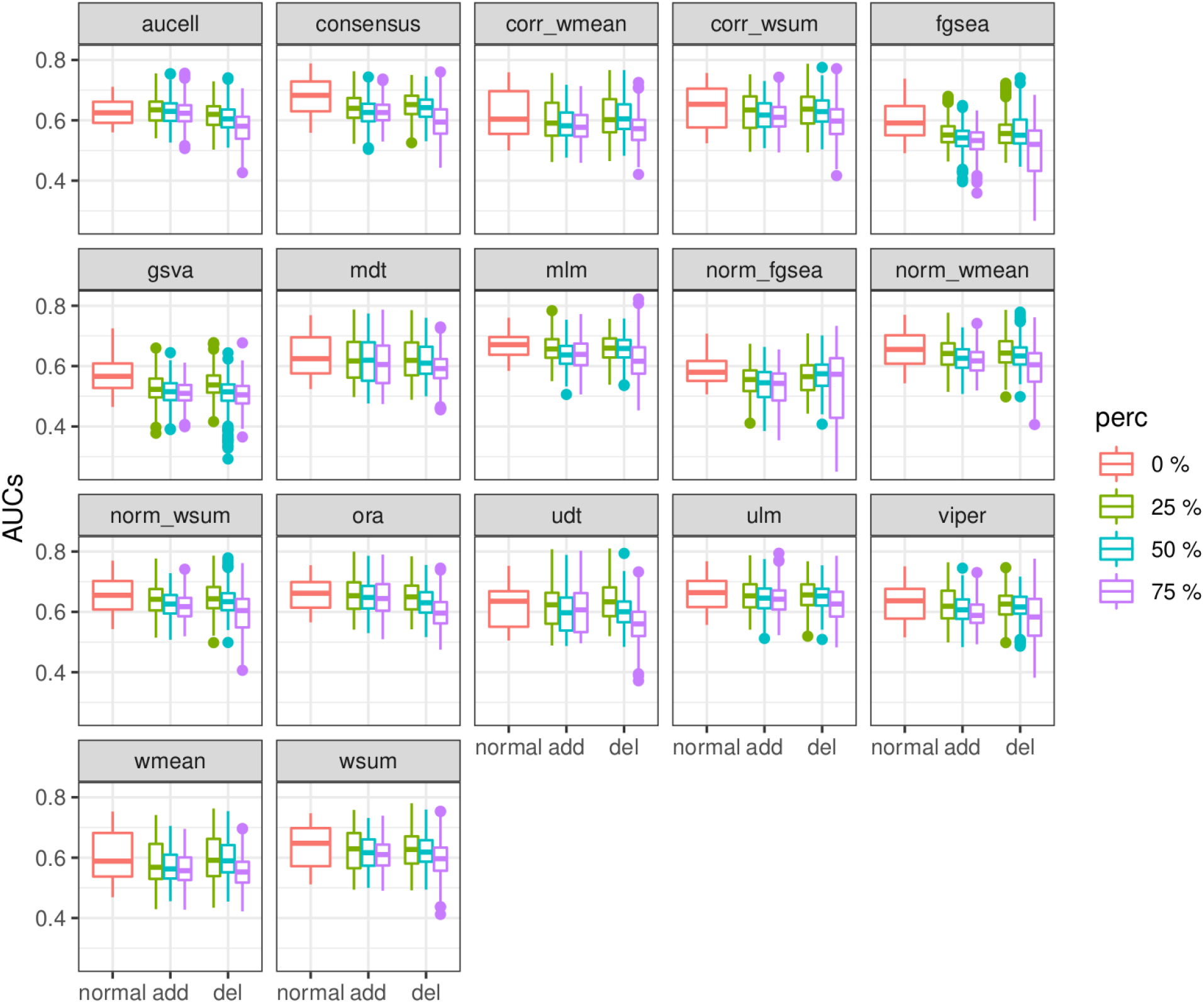
Distributions of AUROCs and AUPRCs for both datasets obtained after adding or deleting edges in the prior knowledge resource.

**Supplementary Figure 7.**
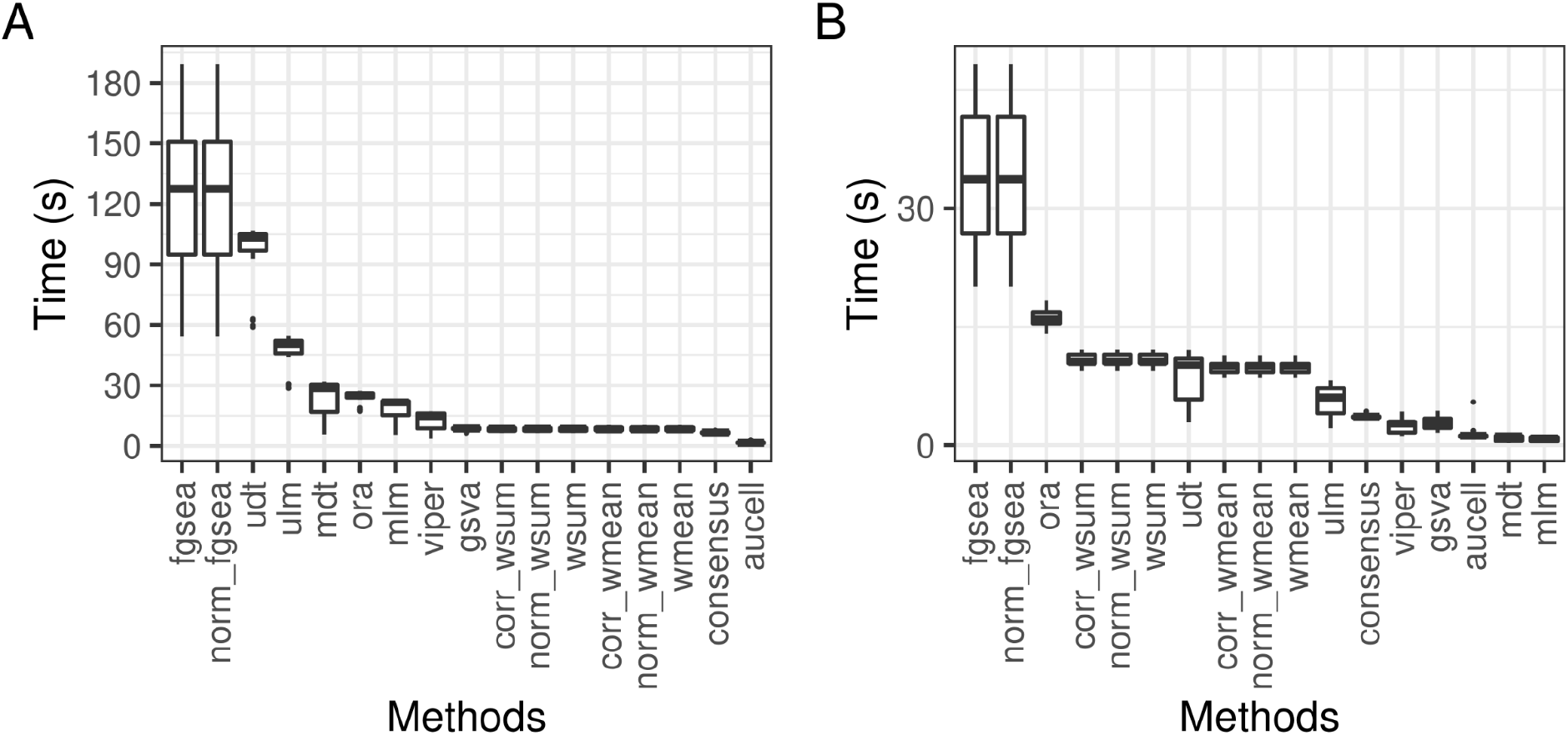
Runtime for each method in the transcriptomics (271 TFs and 92 samples) (A) and phospho-proteomics (59 kinases and 63 samples) (B) datasets. decoupleR was run 31 times in the network robustness experiment (5 permutations, 3 percentages, 2 modes and a normal run) for each dataset in a laptop with an Intel(R) Core(TM) i7-8550U CPU @ 1.80GHz.

## References

Aibar, S. et al. (2017) SCENIC: single-cell regulatory network inference and clustering. Nat. Methods, 14, 1083–1086.

Alhamdoosh, M. et al. (2017) Combining multiple tools outperforms individual methods in gene set enrichment analyses. Bioinformatics, 33, 414–424.

Alvarez, M.J. et al. (2016) Functional characterization of somatic mutations in cancer using network-based inference of protein activity. Nat. Genet., 48, 838–847.

Dugourd, A. and Saez-Rodriguez, J. (2019) Footprint-based functional analysis of multiomic data. Current Opinion in Systems Biology, 15, 82–90.

Fisher, R.A. (1922) On the Interpretation of X2 from Contingency Tables, and the Calculation of P. Journal of the Royal Statistical Society, 85, 87.

Garcia-Alonso, L. et al. (2019) Benchmark and integration of resources for the estimation of human transcription factor activities. Genome Res., 29, 1363–1375.

Geistlinger, L. et al. (2020) Toward a gold standard for benchmarking gene set enrichment analysis. Brief. Bioinformatics.

Hänzelmann, S. et al. (2013) GSVA: gene set variation analysis for microarray and RNA-seq data. BMC Bioinformatics, 14, 7.

Hernandez-Armenta, C. et al. (2017) Benchmarking substrate-based kinase activity inference using phosphoproteomic data. Bioinformatics, 33, 1845–1851.

Holland, C.H. et al. (2020) Robustness and applicability of transcription factor and pathway analysis tools on single-cell RNA-seq data. Genome Biol., 21, 36.

Kolde, R. et al. (2012) Robust rank aggregation for gene list integration and meta-analysis. Bioinformatics, 28, 573–580.

Maciejewski, H. (2014) Gene set analysis methods: statistical models and methodological differences. Brief. Bioinformatics, 15, 504–518.

Mathur, R. et al. (2018) Gene set analysis methods: a systematic comparison. BioData Min., 11, 8.

Ritchie, M.E. et al. (2015) limma powers differential expression analyses for RNA-sequencing and microarray studies. Nucleic Acids Res., 43, e47.

Sergushichev, A. (2016) An algorithm for fast preranked gene set enrichment analysis using cumulative statistic calculation. BioRxiv.

Teschendorff, A.E. and Wang, N. (2020) Improved detection of tumor suppressor events in single-cell RNA-Seq data. BioRxiv.

Therneau, T. and Atkinson, B. (2019) rpart: Recursive Partitioning and Regression Trees.

Väremo, L. et al. (2013) Enriching the gene set analysis of genome-wide data by incorporating directionality of gene expression and combining statistical hypotheses and methods. Nucleic Acids Res., 41, 4378–4391.

Wright, M.N. and Ziegler, A. (2017) ranger : A Fast Implementation of Random Forests for High Dimensional Data inC++ andR. J. Stat. Sofiw., 77, 1–17.

Yilmaz, S. et al. (2021) Robust inference of kinase activity using functional networks. Nat. Commun., 12, 1177.

